# The Phantom of the PCR: detection and consequences of spurious UMIs in mainstream RNA sequencing

**DOI:** 10.64898/2026.08.01.742199

**Authors:** Ken Sugino, Tzumin Lee

## Abstract

Unique molecular identifiers (UMIs) support digital molecular counting by tagging molecules before amplification, but assume that UMIs are incorporated only during reverse transcription. Residual UMI-bearing oligonucleotides can instead reprime during preamplification PCR, creating “phantom” UMIs on genuine cDNA that inflate counts and evade standard deduplication. We model phantom generation as a two-state branching process and show that it produces a heavy-tailed reads-per-UMI distribution distinct from that of true UMIs. Using this signature, PhantomUMI detects and estimates contamination from clone-size distributions, subject to a coverage-dependent identifiability limit. Across 23 datasets spanning published studies and companion experiments, we find signatures consistent with phantom-UMI generation, including in current 10x GEM-X chemistry. Simulations show that phantoms inflate molecule counts and distort fold-changes. Model-based correction removes average count inflation but does not recover the distorted fold-changes, indicating that phantom UMIs are best prevented experimentally, as implemented in the companion Omega-seq method.

---

UMIs have become the standard for digital molecular counting in RNA sequencing: by tagging each cDNA molecule with a random barcode before amplification, reads that share a UMI are collapsed to a single molecule, nominally erasing PCR duplication bias [1, 2]. This guarantee rests on one assumption: that a UMI is attached to a transcript exactly once, during reverse transcription (RT). In practice the oligonucleotides that carry UMIs (the RT primer in tag-based protocols, the template-switching oligo, TSO, in full-length protocols) are not fully consumed by RT and persist into the preamplification PCR. There they can prime synthesis on already-amplified cDNA, attaching a fresh UMI to a genuine transcript. We call the resulting barcodes *phantom UMIs*. The term has been applied before to UMIs created by sequencing or PCR errors, index hopping, and chimeras [3, 4]; because those artifacts arise from rare, random events, they carry markedly fewer reads than true UMIs and can be suppressed by error-aware collapse [5] or a read-count threshold [3]. The phantoms we describe are fundamentally different: they are error-free barcodes on genuine cDNA, and, once created, they are amplified through the remaining PCR cycles. Phantom UMIs therefore evade both error-aware collapse and read-count filtering, and they inflate molecule counts within a single library.

To understand the artifact we model UMI amplification as a two-state branching process (Fig. 1A; Methods). At each cycle a molecule is either primer-terminated (T2 in Fig. 1A: competent for exponential PCR) or TSO/RT-terminated (T1), and residual-oligo priming converts between states at rates proportional to *w*_tso_ (residual-oligo repriming of TSO-ended templates; T1→T1) and *w*_pcr_ (residual-oligo repriming of PCR-primer-ended templates; T2→T1). The UMI-bearing oligo is the TSO in full-length and 5*^′^* droplet protocols and the RT (oligo-dT) primer in 3*^′^* droplet and total-RNA protocols; the scheme is identical either way, and we write *w*_tso_ for its repriming rate throughout. The model makes a sharp prediction: whereas true UMIs, all present from cycle zero, amplify to a narrow clone-size distribution, phantom UMIs are generated continuously across cycles and therefore inherit a heavy-tailed, power-law-like distribution of reads per UMI (derived in Methods). The empirical clone-size (reads-per-UMI) histogram is therefore a direct fingerprint of contamination: a clean library gives a narrow, unimodal distribution set only by PCR amplification noise (a Galton–Watson branching process, approximately negative-binomial), whereas even a modest phantom load adds a characteristic excess of small clones (Fig. 1B).

**Fig. 1.**
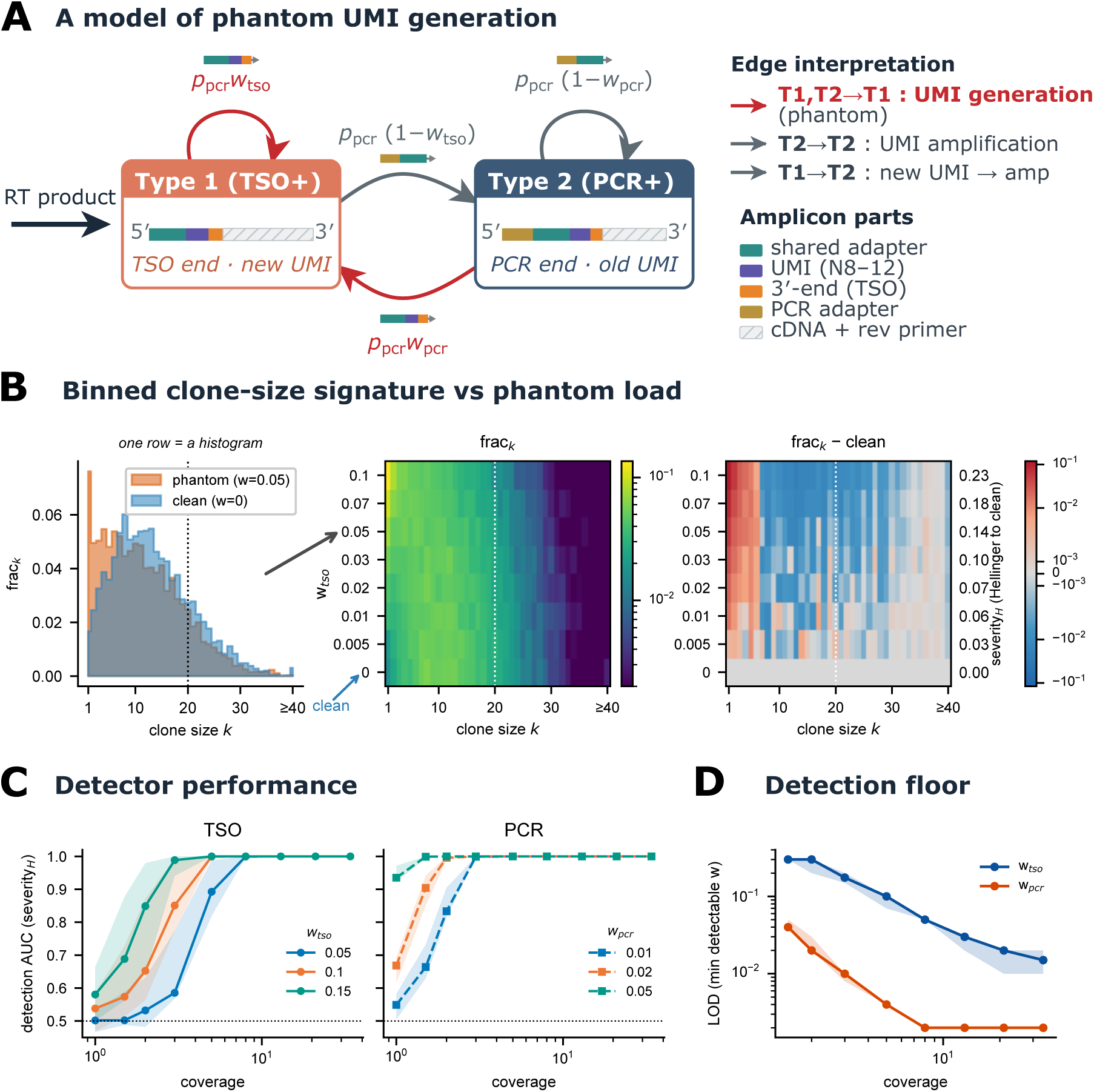
A branching-process model of phantom-UMI generation and its detection. **A**, Two-state model: molecules bearing a terminal TSO (type 1) or a proper PCR primer (type 2) interconvert during PCR; residual-oligo priming (rates ∝ *p*_pcr_*w*_tso_, *p*_pcr_*w*_pcr_) generates new UMIs (Methods). **B**, Binned reads-per-UMI (clone-size) histogram versus phantom load: true UMIs give an approximately negative-binomial signature, phantoms add a heavy-tailed excess of small clones; severity *H* (Hellinger distance to the nearest clean grid cell) quantifies the departure. **C**, Detection AUC of the detector’s severity*_H_* score (Hellinger distance of the binned clone-size histogram to the nearest clean cell, the statistic thresholded by the two-stage gated rule; Methods) versus coverage, for TSO (*w*_tso_, solid) and PCR (*w*_pcr_, dashed) contamination. Line, median across PCR efficiencies *p*_pcr_ ∈ {0.5, 0.6, 0.7, 0.8}; shaded band, full range across those efficiencies. **D**, Detection floor: minimum detectable contamination versus coverage. *w*_pcr_ is detectable ∼10× lower than *w*_tso_; both floors fall with coverage.

A first, coarse diagnostic follows directly from this heavy tail. As sequencing depth *S* increases, the fraction of reads that reveal a new UMI, *Q*(*S*) = *U* (*S*)*/S* (where *U* (*S*) is the number of distinct UMIs recovered at depth *S*), declines with a log–log slope (the logQ-slope) that approaches −1 for a clean library and flattens toward 0 as a phantom tail is added. This slope is the diagnostic used in the companion Omega-seq study [6] and is derived in full in Methods. A single scalar index summarizing the severity of phantom-UMI contamination is very useful; however, its clean baseline is not a fixed constant but drifts with coverage and PCR efficiency: a clean library’s logQ-slope, for example, ranges from near −1 at deep coverage to well above it when reads per UMI are few. Without knowing these parameters, no single global threshold separates clean from contaminated libraries, and shallow clean libraries are misread as phantom-positive (Supplementary Fig. S3). Referencing the full binned clone-size distribution to the nearest clean grid cell at each library’s own coverage and efficiency removes this ambiguity, and mechanistic, distribution-based models are effective for related UMI-counting problems [3].

We exploit this fingerprint to detect and quantify phantoms directly from data. Our PhantomUMI software bins the pooled clone-size histogram and matches it, by Hellinger distance, to the nearest cell of a simulation grid spanning *w*_tso_, *w*_pcr_ and coverage, returning calibrated contamination estimates without any spike-in or external molecular reference [7]. On simulated libraries the detector achieves high discrimination (Fig. 1C): repriming on PCR-primer-ended templates (*w*_pcr_), which seeds the largest clones, is detectable at far lower sequencing coverage than TSO repriming (*w*_tso_), and the minimum detectable contamination falls with coverage for both (Fig. 1D). This floor is reached in the shallowest droplet data (coverage ≈ 1–2 reads per UMI), where true and phantom molecules alike appear as single-read UMIs and cannot be told apart, an identifiability limit: at coverage this low, true and phantom singletons cannot be distinguished from clone-size information alone.

Applying PhantomUMI to 23 single-cell RNA-seq datasets spanning published studies and companion experiments reveals that phantom UMIs are pervasive and dominated by residual-oligo repriming (*w*_tso_) of TSO-ended rather than PCR-primer-ended templates (Fig. 2A). Full-length protocols span an order of magnitude in contamination: run at their published conditions, FLASH-seq [8] is the most affected (*w*_tso_ ≈ 0.4), whereas Smart-seq3 [9] is cleaner, consistent with its forward PCR primer outcompeting residual TSO; explicit post-RT cleanup lowers *w*_tso_ further. The Smart-seq3 and molecular-spike samples that match FLASH-seq in Fig. 2A are the deliberate low-primer arms of those studies’ titrations (0.1 *µ*M forward primer), not their standard protocol, and they place an upper bound on what primer competition can cost. Critically, the artifact is not confined to niche full-length methods. Libraries from the currently dominant droplet chemistry, 10x Genomics GEM-X v4 [10], show a reproducible clone-size signature consistent with repriming by residual UMI-bearing capture primer.

**Fig. 2.**
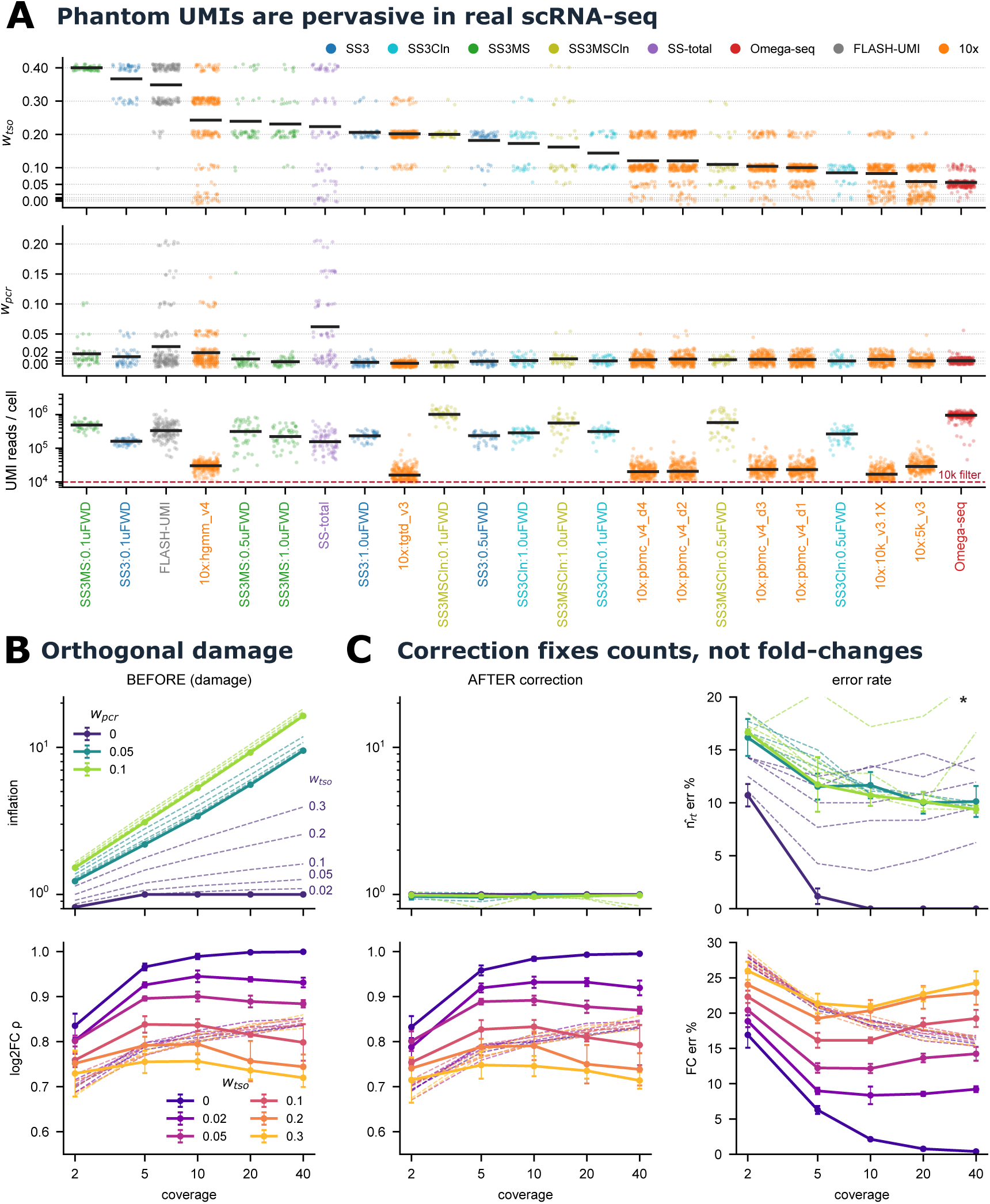
Phantom UMIs are pervasive in real data and their downstream damage is only partly correctable. **A**, Per-cell *w*_tso_, *w*_pcr_ and UMI-reads/cell across 23 datasets (dots, cells; bars, medians), sorted dirtiest to cleanest; phantoms are pervasive and dominated by residual-oligo repriming of TSO-ended templates (*w*_tso_) rather than PCR-primer-ended templates (*w*_pcr_), including in current 10x GEM-X v4 chemistry. Dataset abbreviations, accessions, processing and per-dataset medians are defined in Supplementary Tables B1 and B2; *y*-axis ticks are the discrete simulation-grid values the detector matches to, so the spacing is non-uniform by construction. **B**, Damage versus coverage. Count inflation (top; observed molecules per true molecule, *n̂/n*_rt_; driven by *w*_pcr_) grows with sequencing depth; fold-change rank correlation (bottom) is degraded by *w*_tso_. Colour encodes the base parameter; within each colour the solid line sets the orthogonal parameter to zero and the dashed lines vary it at that fixed base value. **C**, After model-based correction (estimated rates): inflation returns to ∼1× but a ∼10% per-gene residual remains (top), and fold-change correlation is essentially unchanged (bottom); error-rate panels quantify the residuals. ^∗^At *w*_tso_ = 0.2, *w*_pcr_ = 0.1 the two contamination modes become jointly unidentifiable in this shallow, doubly contaminated regime, so a single clean-referenced match cannot separate them and the correction degrades (marked dashed line; Methods).

What do phantoms cost downstream? Because *w*_pcr_ seeds exponentially growing clones (Fig. 1A, T2→T1), it inflates absolute molecule counts, and, counterintuitively, the inflation grows with sequencing depth, as deeper reads resolve ever-smaller phantom clones (Fig. 2B). TSO repriming, by contrast, mainly corrupts relative quantities due to stochastic amplification of the generated phantom UMIs, degrading the rank correlation of fold-changes between conditions. We asked whether these harms can be reversed computationally. Established UMI pipelines collapse sequencing errors [5] and correct for molecules *lost* to amplification or sequencing inefficiency [3], but the phantom problem is the opposite, a spurious gain of molecules. A correction that estimates the contamination rates from each library and inverts the modelled inflation (Methods; Fig. B6) removes the count bias almost completely, driving inflation back to ∼1× (Fig. 2C). But it leaves two irreducible problems. First, a residual per-gene error of ∼10% persists: the correction restores counts on average (the median inflation curve), but individual genes scatter around it owing to finite-sampling (shot) noise. Second, and more fundamentally, fold-changes are not restored: because the correction rescales counts almost globally (nearly the same factor for every gene) it preserves their ratios, so the fold-change distortion survives essentially unchanged (Fig. 2C). Computational correction is therefore a diagnostic-grade mitigation, not a cure.

These results reframe phantom UMIs as a systematic, quantifiable, and, for differential analysis, largely uncorrectable bias in mainstream UMI sequencing. The practical consequence is twofold. First, phantom contamination should be measured and reported as a routine quality-control metric; the PhantomUMI package does this from the clone-size distribution alone, requiring nothing beyond the reads already in hand. Second, because the damage to fold-changes cannot be undone after the fact, phantom UMIs must be prevented at the bench. In the companion study we introduce Omega-seq [6], a full-length library-preparation method that prevents phantom generation at its source, by digesting the uracil-bearing TSO, using a uracil-intolerant polymerase that stalls at template uracil and cannot amplify residual TSO-derived strands, and raising the ISPCR primer concentration to outcompete residual priming. Detection and prevention are complementary: PhantomUMI reveals the phantom in existing and new data, and Omega-seq banishes it.

## Declarations

## Ethics approval and consent to participate

Not applicable

## Consent for publication

Not applicable

## Data availability

All datasets and their public accessions are listed in Supplementary Table B1.

## Code availability

All analysis code and the “PhantomUMI” Python package are available on GitHub at [XXX].

## Supplementary information

Appendix B (Supplementary Tables and Supplementary Figures) is available with the online version of this article. Appendix A is the Methods of this article, including the derivations underlying the model and the detector, and is part of the main, peer-reviewed text rather than supplementary material.

## Author contributions

K.S. developed the mathematical framework for phantom UMI generation, analyzed the data, developed the software package, and wrote the manuscript. T.L. secured funding, supervised the project, and contributed to intellectual discussions from project conception through manuscript completion.

## Appendix A Methods

### A.1 Mathematics of Phantom UMI generation

Let *T_k_* and *P_k_* denote the number of molecules of type 1 (bearing a terminal TSO) and type 2 (bearing a proper PCR primer) at the *k*-th PCR cycle, respectively. The transition probabilities between states are determined by the overall PCR efficiency (*p_pcr_*), the proportion of type 1 molecules generating new type 1 molecules (*w_tso_*), and the proportion of type 2 molecules generating new type 1 molecules (*w_pcr_*). We define the complementary transition weights as 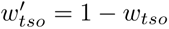 and 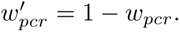

The system’s Markov dynamics can be described by the linear recurrence relation:

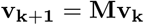

where

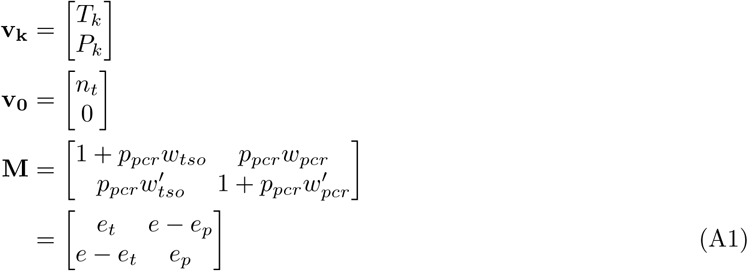

Here, *n_t_* is the initial number of true (RT-derived) transcript molecules of a single gene; the simulator and data-analysis sections write the same quantity as *n*_rt_ (§B.1, §B.5). The amplification rates are defined as:

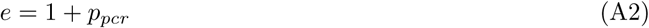

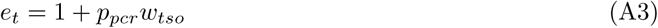

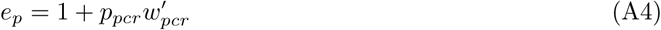

corresponding to the total amplification rate (*e*), the type 1 branch amplification rate (*e_t_*), and the type 2 branch amplification rate (*e_p_*).

The transition matrix **M** yields the following eigenvalues:

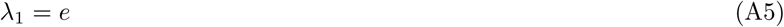

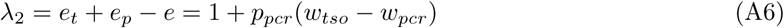

and corresponding eigenvectors:

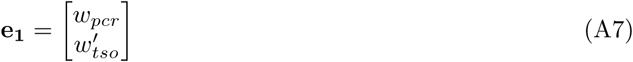

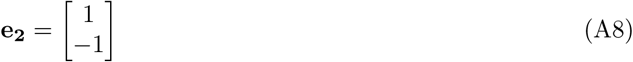

Note that *λ*_1_ governs the total PCR efficiency, while *λ*_2_ governs the efficiency of the differential generation of type 1 copies from type 1 versus type 2 branches. The state vector at cycle *k* is given by:

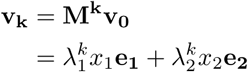

where the initial condition vector **v_0_** is expressed as:

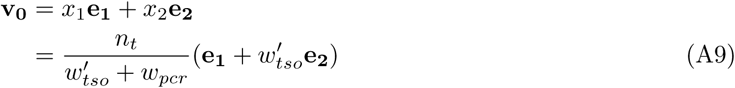

Solving for the components yields:

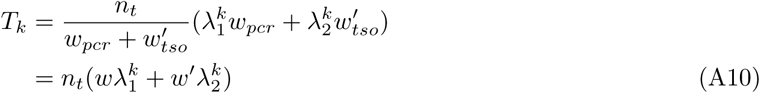

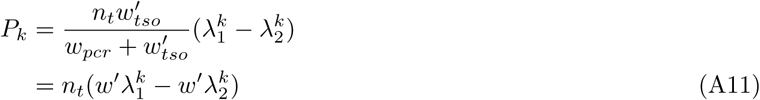

where the scaling weights are:

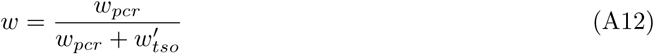

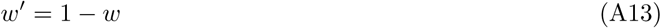

Equations A10 and A11 demonstrate that the exponential growth of both *T_k_*and *P_k_* is primarily driven by the dominant eigenvalue *λ*_1_. This growth is partitioned by a fixed ratio (*w* to *w^′^*) determined by the transition rates between type 1 and type 2, and is modified by the continuous transfer of molecules between the branches 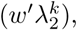 ultimately yielding a total molecule count 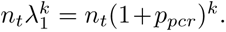 The number of new UMIs generated specifically at cycle *k* (for *k >* 0) is:

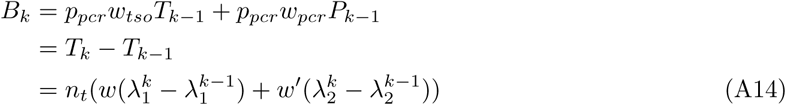

We set *B*_0_ = *n_t_* to represent the initial pool of true (non-phantom) UMIs. Then the cumulative number of UMI families generated up to cycle *k*, *S_k_*, is:

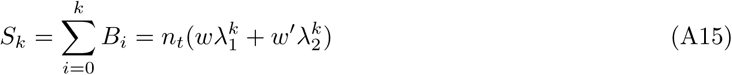

Because UMIs generated at earlier cycles have longer to amplify, the condition of being generated at or before cycle *k* is equivalent to achieving a final clone size of at least *X_k_*. Therefore, the complementary cumulative distribution function (CCDF) of the UMI clone sizes is:

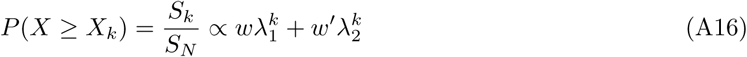

where the variable *X_k_*represents the expected final clone size of a UMI family originating at cycle *k*, and *S_N_* is the total number of UMI families generated across all *N* cycles.

Each new UMI generated in the type 1 branch remains a single copy within that specific branch, as any further amplification of a type 1 molecule creates a distinct UMI. However, at each subsequent cycle, this molecule acts as a template, generating type 2 copies at a rate of 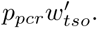 Once in the type 2 branch, these descendants amplify exponentially at a rate of 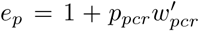 for the remainder of the PCR process. Assuming no sequence collisions between newly generated UMIs, the expected final clone size *X_k_* of a UMI family born at cycle *k* after *N* total cycles is:

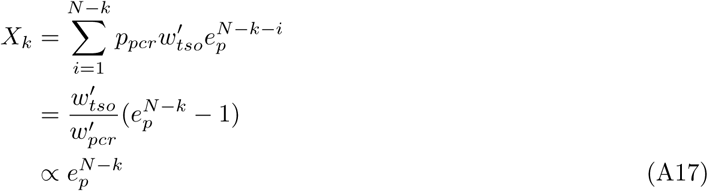

**Fig. A1.**
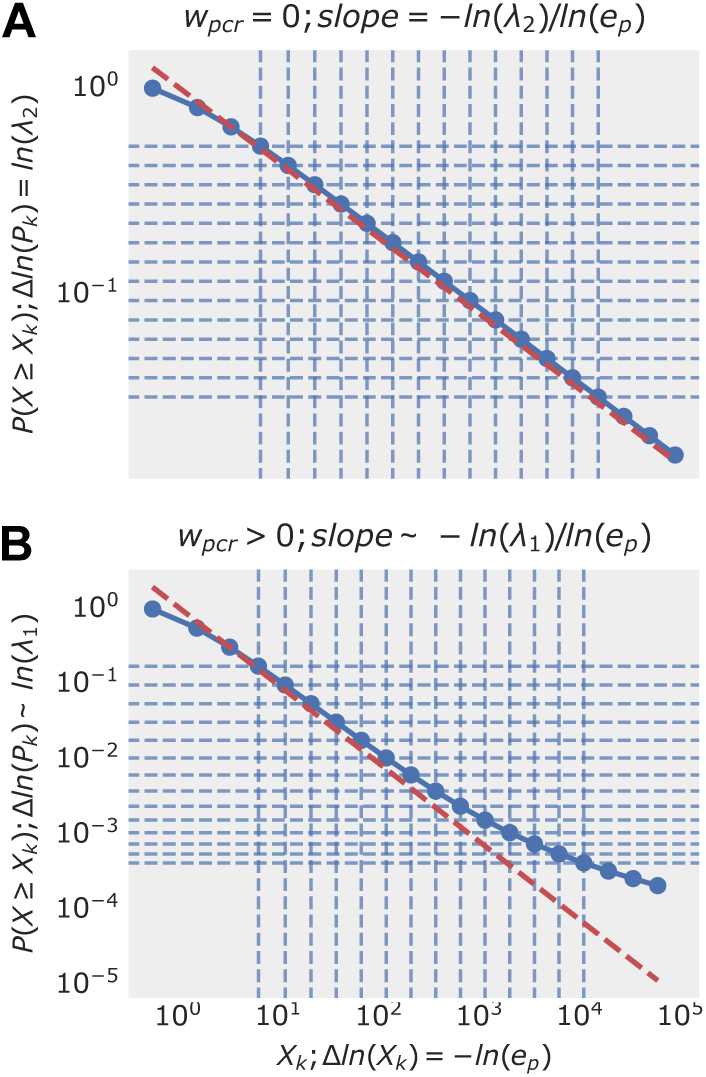
Complementary cumulative distribution function (CCDF) of theoretical UMIs with phantom generation. The x-axis represents the UMI clone size and the y-axis is *P* (*X* ≥ *x*). Parameters: *p_pcr_* = 0.8, *n_t_* = 100, *N* = 20, *w_tso_* = 0.3, with *w_pcr_* = 0 (Panel A) and *w_pcr_* = 0.05 (Panel B). Note that in Panel B, Δ ln(*S_k_*) ∼ ln(*λ*_1_) holds true only for large *k* (i.e., smaller clone sizes *X_k_*).

An example of this distribution in log-log space is shown in Fig. A1, which displays a strict linear relationship (excluding terminal points), characteristic of a power-law distribution. The power-law exponent, the slope in log-log space, can be calculated by taking the ratio of the discrete differences of *S_k_* and *X_k_* in logarithmic space:

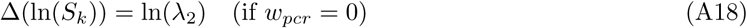

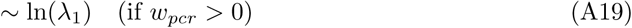

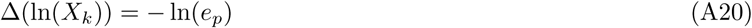

For the regime where *w_pcr_ >* 0, we assume *k* is sufficiently large such that (*λ*_2_*/λ*_1_)*^k^* ≪ 1. Therefore, the exponent of the power-law distribution *P* (*X* ≥ *x*) ∝ *x^−β^* is given by:

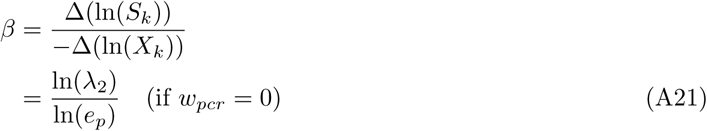

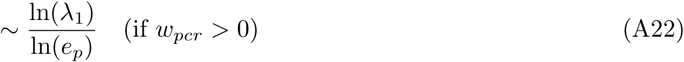

### A.2 Phantom UMIs in the Saturation Curve

The expected probability of detecting a specific unique molecule *UMI_i_* at a sequencing depth of *S* follows the standard Poisson capture probability:

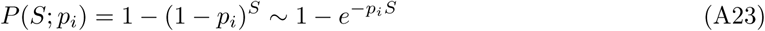

where *p_i_* is the relative frequency of *UMI_i_* in the final amplified library. If a UMI is generated at cycle *k*, it has an expected clone size of *X_k_*, and there are *B_k_* such UMIs. Therefore, its frequency is:

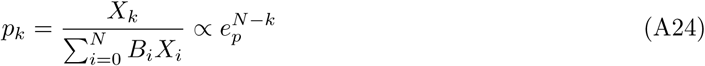

Accordingly, the expected total number of unique UMIs discovered at sequencing depth *S* is the sum of these probabilities:

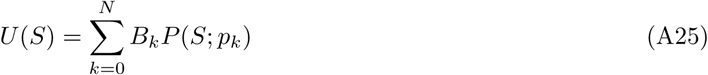

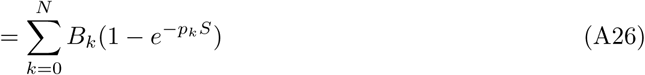

In log-log space, the unit Poisson detection probability curve for a single family transitions asymptotically from a slope of 1 to a slope of 0 at the critical point *S* = 1*/p_k_* (see Fig. A2). Since 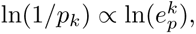 the horizontal spacing between them is exactly ln(*e_p_*). Because each term in Eq. A26 is scaled by *B_k_*, which corresponds to a vertical translation of ln(*B_k_*) in log-log space, the vertical spacing between individual curves is Δ ln(*B_k_*) = ln(*λ*_2_) (if *w_pcr_* = 0) or Δ ln(*B_k_*) ∼ ln(*λ*_1_) (if *w_pcr_ >* 0).

As shown in Fig. A3A (where *w_pcr_* = 0), the individual Poisson detection curves for phantom UMIs (*k >* 0, graded red lines) sequentially accumulate alongside the “true” UMIs (*k* = 0, black dashed line) to form the total discovery curve *U* (*S*) (solid red line). While this summation occurs in linear space, under certain parameter regimes (e.g., Fig. A3A), a specific cycle *k* strictly dominates the discovery rate at any given depth *S*. Consequently, the overall shape of *U* (*S*) in log-log space is tightly approximated by the mathematical envelope of these individual phantom curves. In such cases, the slope of the envelope is ln(*λ*_2_)*/* ln(*e_p_*) = *β* between the boundaries of 1*/p*_0_ and 1*/p_N_*. The necessary condition for this envelope-dominated regime is that the vertical spacing (ln(*λ*_2_)) is much smaller than the horizontal spacing (ln(*e_p_*)), meaning *β* ≪ 1.

Conversely, when *β* ≫ 1 such that Δ ln(*B_k_*) *>* ln(*e_p_*) for all *k >* 0 (Fig. A3B), the massive reservoir of singleton UMIs generated at the final cycle (*k* = *N*) overwhelmingly dominates the earlier cycles, driving the aggregate slope of *U* (*S*) very close to 1.

In the intermediate regime where *β* ∼ 1, the initial linear segments of the individual Poisson detection curves overlap extensively (Fig. A3C), preventing any single cycle *k* from dominating. At *S* = 1*/p*_1_, phantom UMIs from multiple cycles contribute equally; however, as sequencing depth increases to *S* = 1*/p*_2_, the discovery contribution from *k* = 1 saturates. This continuous, cascading saturation causes the slope of *U* (*S*) to become shallower than the theoretical envelope.

**Fig. A2.**
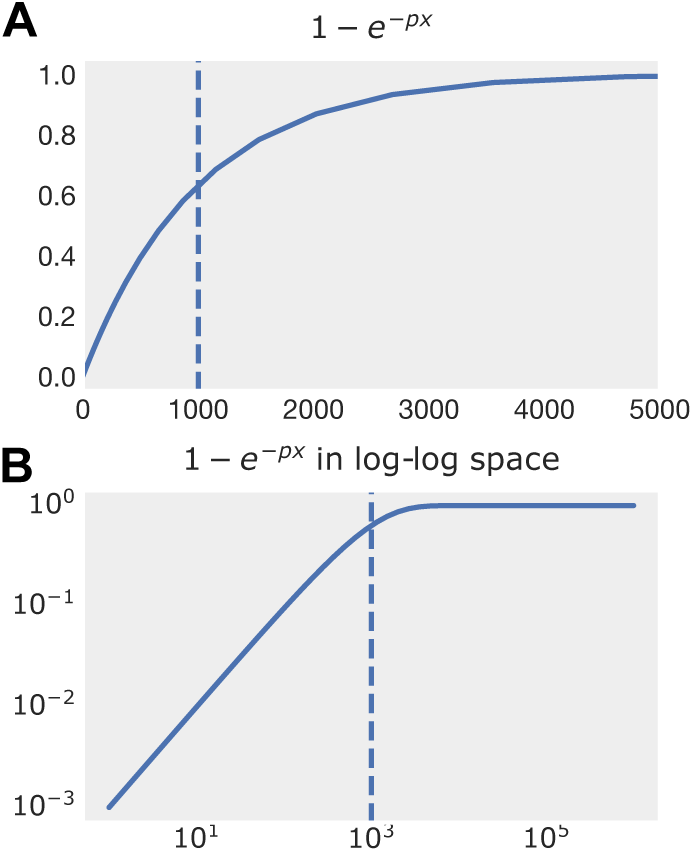
The single Poisson detection probability function 1 − *e*^−*px*^ plotted in linear scale (Panel A) and log-log scale (Panel B). Vertical dashed lines indicate the critical transition point where *x* = 1*/p*. In log-log space, this function can be approximated as two asymptotes (one with a slope of 1, the other with a slope of 0) intersecting at the transition point. Slopes throughout this section refer to the discovery curve *U* (*S*), which flattens to slope 0 once a library saturates; the main text instead quotes slopes of *Q*(*S*) = *U* (*S*)*/S*, which are smaller by exactly one, so the same saturated library approaches −1 there.

Ultimately, the deviation from standard saturation (black dashed line) occurs strictly between the boundaries of 1*/p*_0_ and 1*/p_N_*. The lower bound, 1*/p*_0_, represents the inverse frequency of the true UMIs (roughly the ratio of the total library size to the largest clone size). Because the final phantom UMIs generated at cycle *N* are singletons, the upper bound 1*/p_N_* precisely equals the total number of molecules in the library after PCR. Assuming that sequencing represents multinomial sampling from this final amplified pool, standard artifact-free libraries should flatline (slope = 0) once *S >* 1*/p*_0_, which can be empirically approximated as the ratio of the total sequencing depth to the largest observed UMI clone size. Therefore, a non-zero slope beyond this threshold serves as a mathematical indicator for the presence of phantom UMIs.

### A.3 Multi-gene Aggregation and Gene Detection Saturation

The theoretical framework established above models the phantom UMI dynamics for a single initial transcript family. However, this mathematical behavior scales to an aggregated, multi-gene library. As shown in eq. A14, the number of phantom UMIs generated at cycle *k*, *B_k_*, is strictly proportional to the initial number of true transcripts, *n_t_*. For a transcriptome consisting of G genes, each with an initial transcript count *n_t,g_*, the total number of UMIs originating at cycle *k* is simply the linear sum 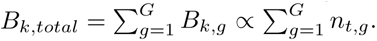 Since the dependency of *B_k,total_* on the cycle *k* remains identical to the single-gene case, differing only by this global scaling factor, the vertical spacing between consecutive cycles, Δ ln(*B_k_*), remains strictly unchanged in log-log space.

Because the expected clone size *X_k_* and the resulting relative frequency *p_k_* depend exclusively on the amplification kinetics (*e_p_*, *w_tso_*, *w_pcr_*) and the remaining PCR cycles (*N* − *k*), they are independent of the initial transcript abundance *n_t_*. Consequently, aggregating across the transcriptome only shifts the absolute vertical position of the unit Poisson curves. The horizontal transition points (1*/p_k_*), the horizontal spacing (ln(*e_p_*)), and the aforementioned vertical spacing (Δ ln(*B_k_*)) remain universally invariant. Thus, the macroscopic geometry of the total discovery curve *U* (*S*) and the power-law exponent *β* are preserved in a full-library context.

**Fig. A3.**
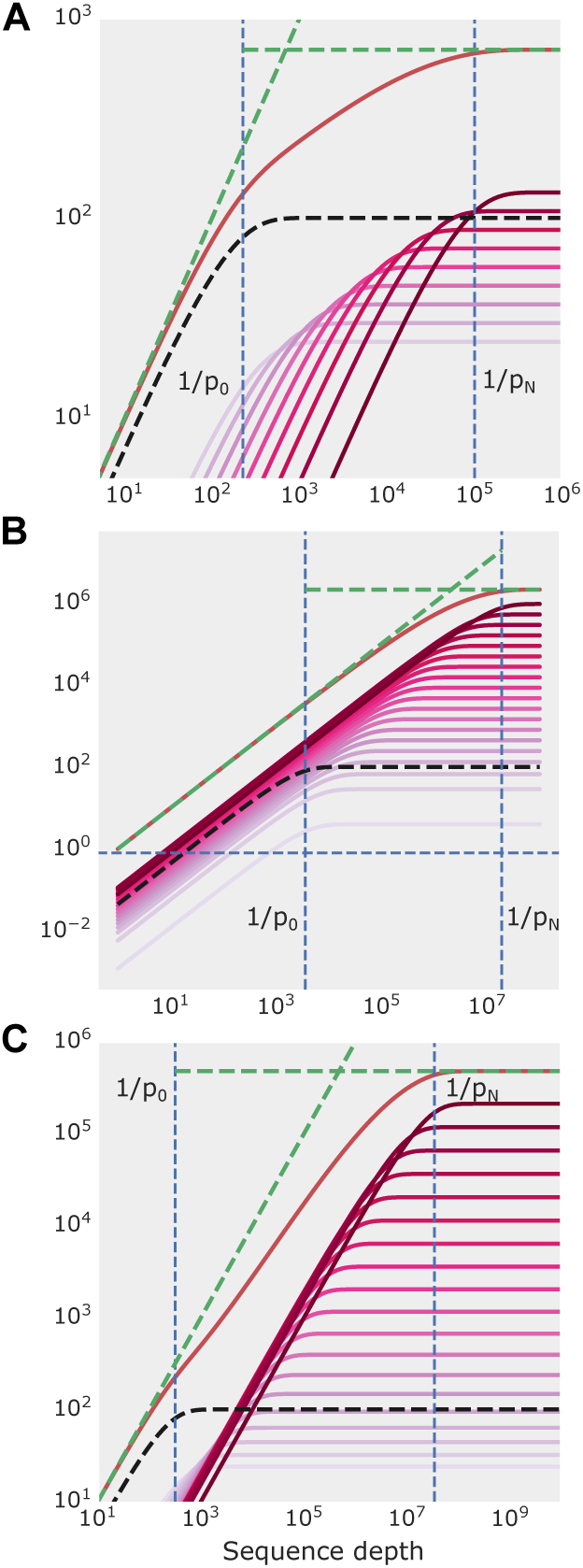
Green dashed lines indicate slopes of 1 and 0. Vertical blue dashed lines mark 1*/p*_0_ (the inverse frequency of a “true” UMI) and 1*/p_N_* (the total library size, where *k* = *N* phantom UMIs are singletons). The black dashed curve indicates the contribution from true UMIs, the graded red curves indicate contributions from phantom UMIs born at subsequent cycles, and the solid red curve represents the total discovery curve *U* (*S*). Parameters: **A** *w_tso_* = 0.3*, w_pcr_* = 0*, N* = 10*, p_pcr_* = 0.8*, n_t_* = 100 **B** *w_tso_* = 0.05*, w_pcr_* = 0.4*, N* = 20*, p_pcr_* = 0.8*, n_t_* = 100 **C** *w_tso_* = 0.3*, w_pcr_* = 0.05*, N* = 20*, p_pcr_* = 0.8*, n_t_* = 100

The same invariance extends to the per-gene form of the metric, in which *Q* is taken as the median across genes of the per-gene ratio of distinct UMIs to reads rather than as the library-level ratio *U* (*S*)*/S* (as in the companion study [6]). Because the amplification kinetics are independent of abundance, every gene follows the same function of its *own* read count, *Q_g_*(*S*) = *Q̃*(*f_g_S*), where *f_g_* is the gene’s share of library reads; in log-log space the per-gene curves are therefore horizontal translates of one another. Since *Q̃* is monotone in its argument, taking the median across genes commutes with it, median*_g_ Q̃*(*f_g_S*) = *Q̃*(*f*_med_*S*) with *f*_med_ the median read share, so the median-of-per-gene-ratios statistic has exactly the same log-log slope as the aggregate curve, evaluated at a depth axis rescaled by *f*_med_. The two definitions therefore share the theory developed here; because the rescaling shifts where a library sits on the common curve, comparisons between libraries must be made at matched coverage (§C.3).

Crucially, this aggregated UMI saturation curve can be directly anchored to the standard gene detection saturation curve. For any given gene, its individual detection probability curve transitions asymptotically at a sequencing depth equal to the inverse of the gene’s total relative frequency in the library (i.e., the library size divided by the total sum of all UMIs belonging to that gene). The theoretical upper bound for this transition point on the x-axis is defined by the rarest possible transcript: a single-copy gene (*n_t_*= 1).

**Fig. A4.**
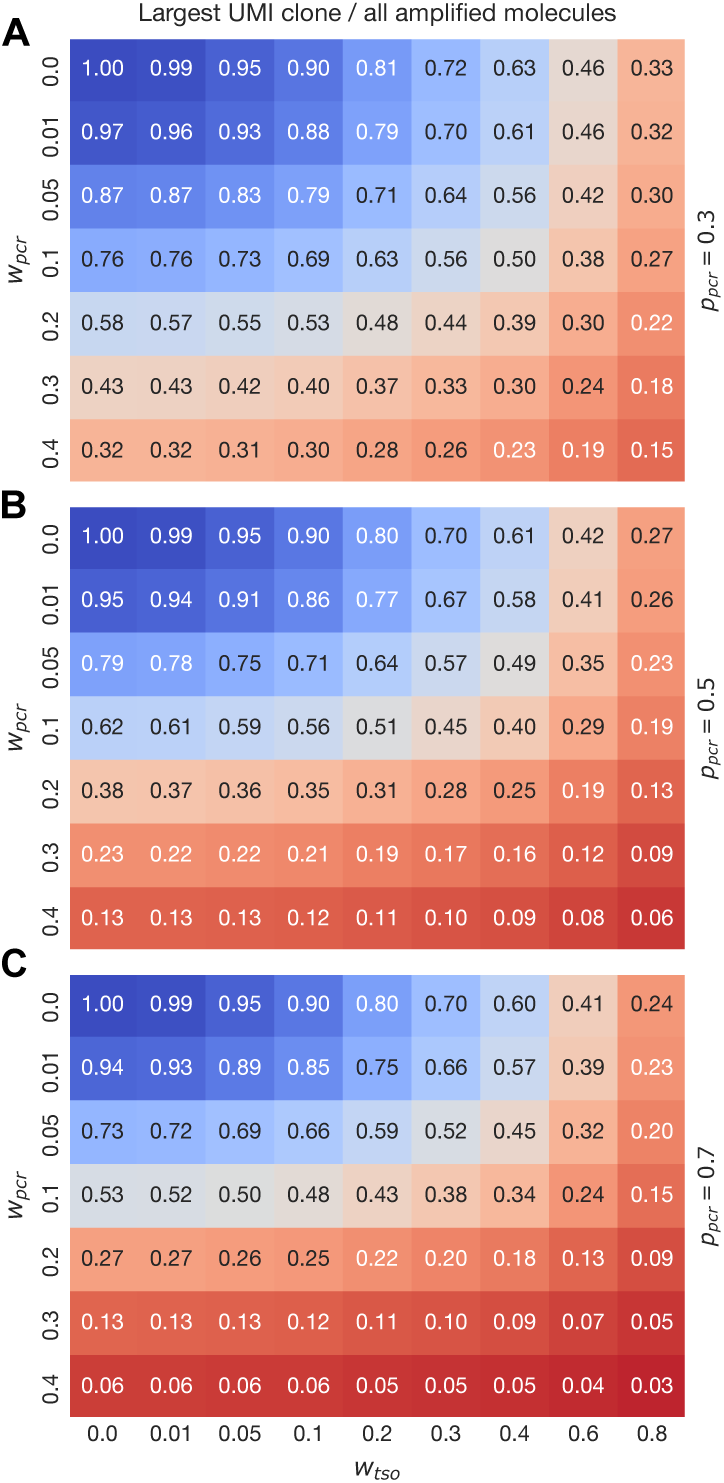
Fraction of amplified molecules belonging to the largest UMI clone. Parameters: initial transcript count *n_t_* = 1, total PCR cycles *N* = 20. Other specific parameters are indicated within the figure. The three panels demonstrate varying base PCR efficiencies (*p_pcr_*). As long as the phantom generation rates remain within practical ranges (*w_pcr_* ≤ 0.05 and *w_tso_* ≤ 0.4), the largest UMI clone contains a substantial fraction of all amplified molecules derived from the initial transcript.

Within the practical parameter ranges of *w_tso_* and *w_pcr_* (See Fig. A4), the amplification dynamics dictate that the largest UMI clone contains at least 50% of all amplified molecules derived from that initial transcript. Mathematically, this means the relative frequency of the largest unique UMI (*p*_0_) and the relative frequency of the entire single-copy gene (*P_single copy_*) are on the same order of magnitude, making their transition points nearly identical (1*/p*_0_ ∼ 1*/P_single copy_*).

Because the single-copy gene detection point represents the maximum possible transition point for any gene in the library, the global gene detection saturation curve will flatten out once the sequencing depth surpasses this threshold. Therefore, the flattening of the standard gene detection curve serves as an empirical indication that the unique UMI saturation curve has entered the relevant analytical region (*S >* 1*/p*_0_). Under the model, continued growth of the UMI-discovery curve beyond this boundary is driven by the sequential discovery of phantom UMIs.

**Fig. B5.**
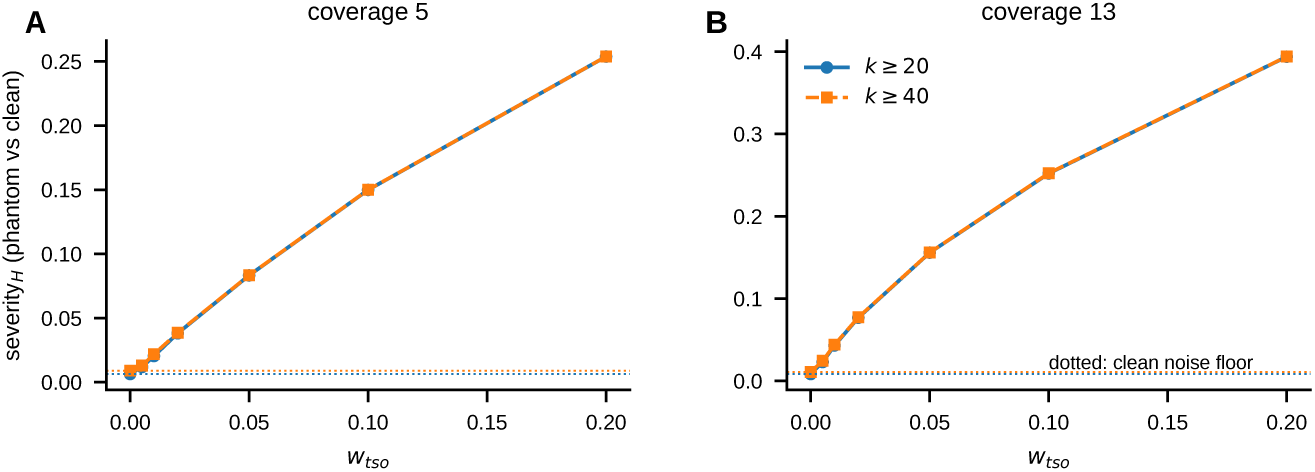
Clone-size binning (*k* ≥ 20 vs *k* ≥ 40). Phantom-versus-clean severity*_H_* versus *w*_tso_ at coverage 5 (A) and 13 (B): the two binnings are nearly identical; dotted lines mark the clean noise floor.

## Appendix B Supplementary Tables

### B.1 Phantom-UMI simulator

Libraries were generated with a custom Rust simulator (phantom sim). For each of *G* genes, *n*_rt_ true molecules are captured by reverse transcription (binomial capture of the transcript pool at rate *p*_rt_) and each is assigned a UMI drawn uniformly from a space of 4*^L^* sequences (*L* = 8, i.e. 65,536 codes, in all simulations). The pool is amplified for *N* PCR cycles under the two-state branching process described above: at each cycle every molecule is copied with probability *p*_pcr_, and residual-oligo priming mints new UMIs from type-1 and type-2 templates at rates proportional to *p*_pcr_*w*_tso_ (T1→T1) and *p*_pcr_*w*_pcr_ (T2→T1), with UMI collisions resolved within the finite UMI space. The amplified pool is sequenced by multinomial sampling to a chosen read depth, and reads are grouped by (gene, UMI) to give the observed clone sizes (reads per UMI). Unless stated otherwise *N* = 20. Coverage is reported as reads per true molecule (total reads divided by ∑*_g_ n*_rt*,g*_), which is independent of phantom load; the phantom-inflated alternative (reads per *detected* UMI) understates it.

### B.2 The PhantomUMI detector

PhantomUMI operates on the pooled clone-size (reads-per-UMI) histogram of a library, binned into *k* = 1*, …,* 19 with a *k* ≥ 20 catch-all (§B.3). The histogram is compared, by Hellinger distance on the binned frequencies, against a precomputed grid of simulated signatures spanning *w*_tso_, *w*_pcr_, PCR efficiency *p*_pcr_ and coverage, together with a matched grid of clean (*w* = 0) signatures.

Detection uses a two-stage gated rule designed conservatively to minimize false-positive calls. Stage 1 tests whether the observed histogram is compatible with nearby clean grid cells by multinomial resampling. A library is called contaminated if it either strongly rejects the clean null (*p < α*_inner_ = 0.01), or shows both weaker evidence against the null (*p < α*_outer_ = 0.30) and a substantial model-based departure from the matched clean distribution (severity*_H_* ≥ 0.15). The thresholds were selected to maintain an empirical false-positive rate near zero across clean validation simulations while retaining sensitivity to detectable contamination. The detection AUC in Fig. 1C is the ROC-AUC of the continuous severity*_H_* statistic that this rule thresholds; the detection floor in Fig. 1D is the smallest contamination *w* recovered at a false-negative rate ≤ 10% and 1% false-positive rate.

### B.3 Choice of clone-size binning

The detector bins the clone-size histogram into *k* = 1*, …,* 19 with a *k* ≥ 20 catch-all. Extending the catch-all to *k* ≥ 40 increases the phantom-versus-clean Hellinger separation only marginally, and the extra bins also raise the clean-versus-clean noise floor, so the net gain is negligible across the detection regime (Fig. B5). The *k* ≥ 20 binning is therefore retained.

**Table B1.**
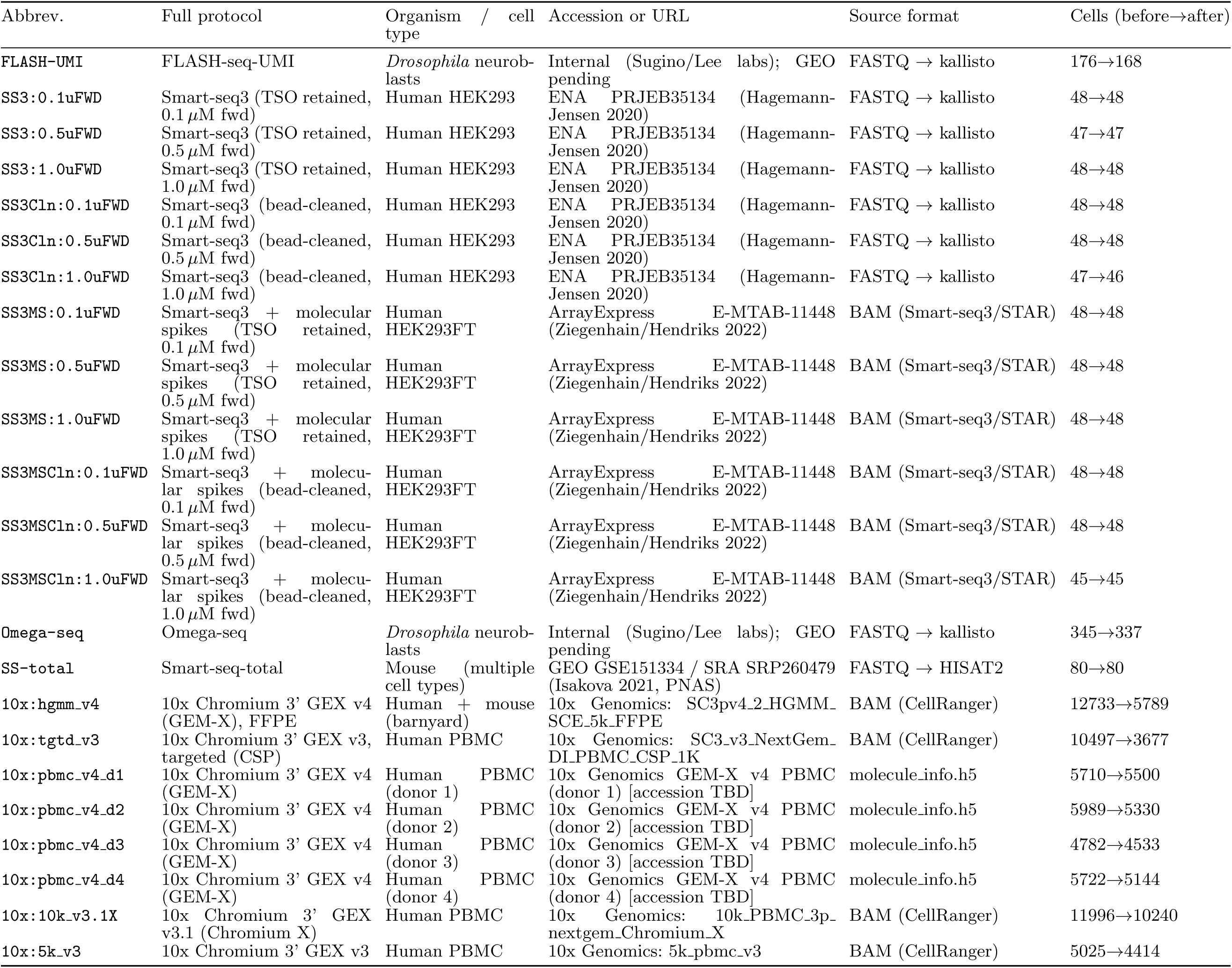
Data sources used in the phantom-UMI benchmark (provenance). The 23 datasets of Fig. 2A; abbreviations are the panel labels. Cells: before → after the ≥ 10^4^ UMI-read credibility filter (reads per cell = summed clone sizes), which removes empty/ambient droplets and failed wells; the detector was run on every cell passing it, and all per-cell estimates and medians reported here and in Fig. 2A are computed over those cells. For the four GEM-X v4 PBMC donors the before-count is the CellRanger pass-filter set of the molecule-info file (per-donor breakdown in Supplementary Fig. S1). Technical parameters and estimated contamination are in Supplementary Table B2.

**Table B2.**
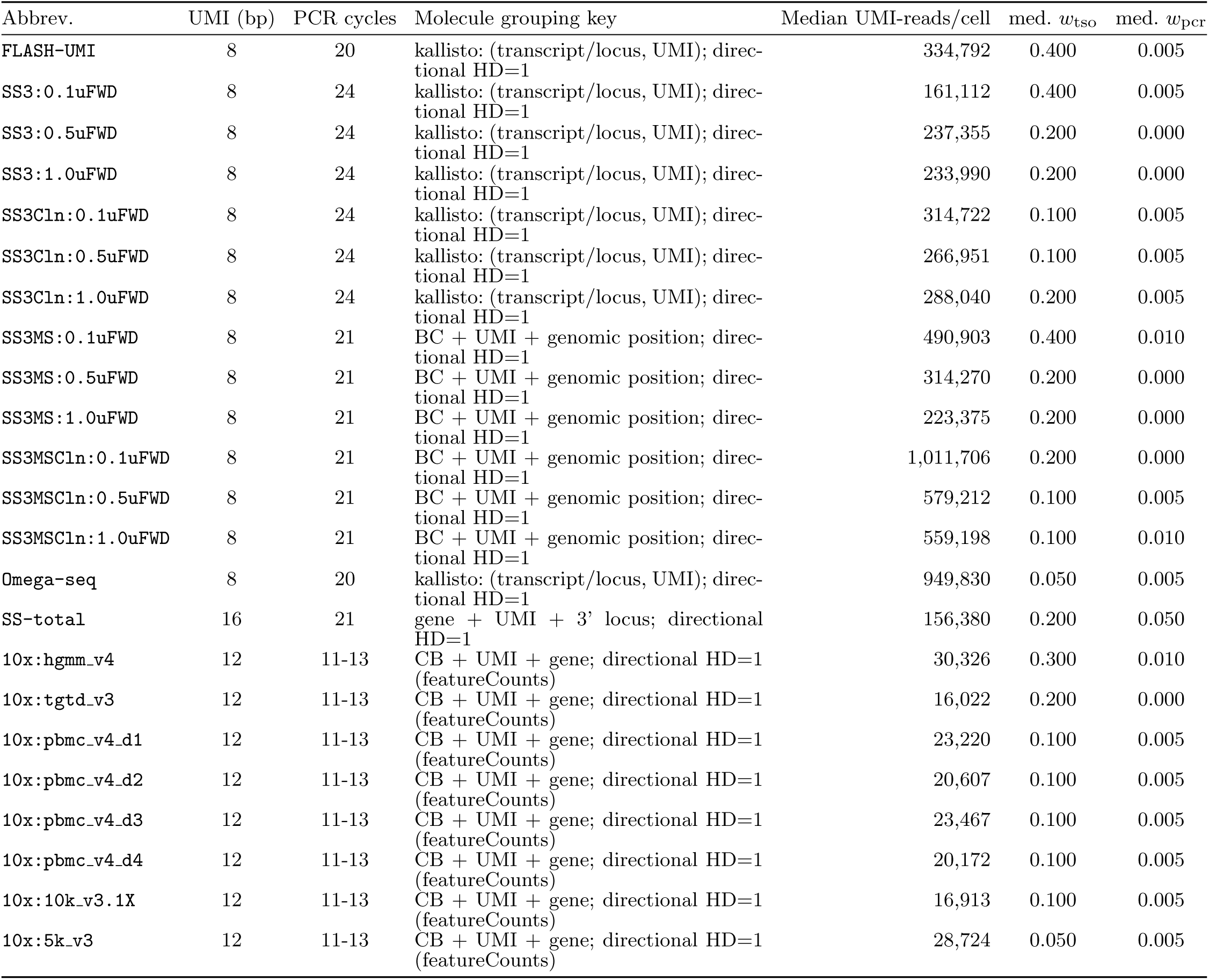
Data sources (technical parameters and estimated contamination). Companion to Supplementary Table B1. Per-cell contamination was estimated with PhantomUMI (detect severity gated) against the v4b lookup grid, except Omega-seq, which was matched against a grid simulating its sequencing of both TSO-ended and PCR-primer-ended molecules; medians over the cells passing the ≥ 10^4^ UMI-read filter. Coverage is summarized by the median UMI-bearing reads per cell (phantom-robust depth; reads per UMI is not used, as its denominator is inflated by phantoms). Molecule grouping is the key on which reads were collapsed to a UMI (directional HD=1). PCR-cycle counts are the preamplification cycles reported in each protocol’s Methods; for 10x they are the cDNA-amplification cycles of the vendor user guide, which are set per run from the targeted cell recovery (*<*500 cells, 13; 500–6,000, 12; *>*6,000, 11) and are identical across the v3, v3.1 and GEM-X v4 kits used here, so the range rather than the per-run value is given. Library index-PCR cycles are excluded throughout, as they cannot create new UMIs.

**Table B3.**
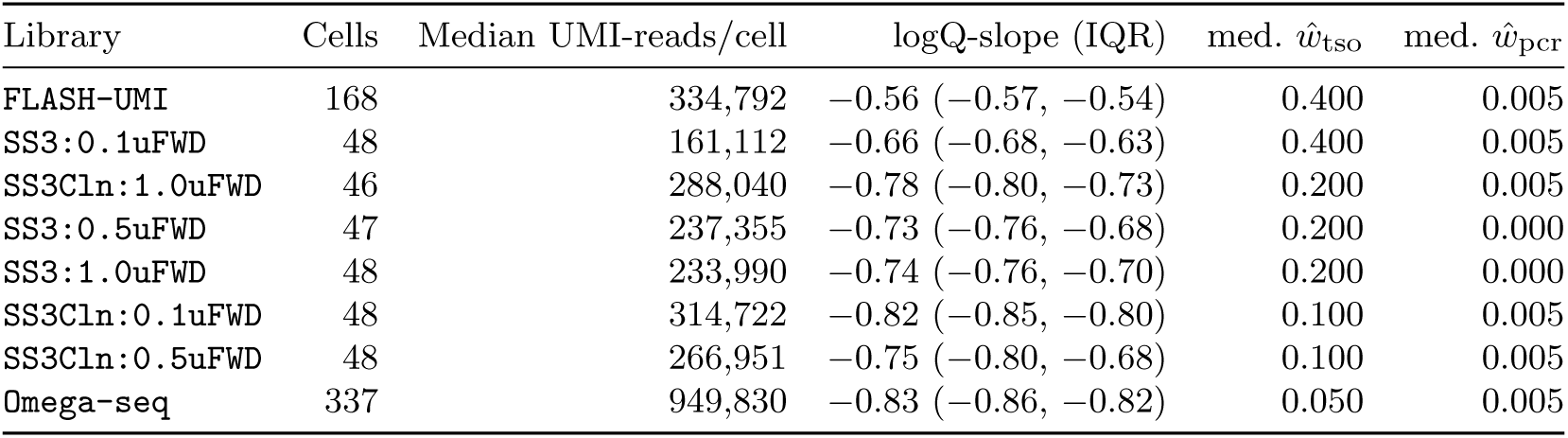
The logQ-slope and the PhantomUMI estimates agree on the libraries scored by both studies. Smart-seq3 at three forward-primer concentrations with and without pre-PCR bead cleanup, FLASH-seq-UMI and Omega-seq, ordered by *ŵ*_tso_. The logQ-slope is the library-level form *Q*(*S*) = *U* (*S*)*/S* computed per cell from the same collapsed clone sizes the detector consumes (median, with interquartile range); the companion study [6] measures the median-of-per-gene form on these libraries with an independent pipeline, which shares the same log-log slope (§A.3) and reproduces this ordering (its Fig. 4F, where Omega-seq scores ≈ −0.8 and FLASH-seq-UMI ≈ −0.3). Absolute values are not comparable between the two forms of *Q* and the two pipelines; the ordering and the separation are. Shallower (less negative) slopes accompany higher *ŵ*_tso_ across the whole panel: Spearman *ρ* = 0.90 between the two columns. The two statistics summarize the same clone-size data in different ways, so this is a consistency check between metrics rather than between independent measurements. All eight are deep full-length libraries (≥ 1.6 × 10^5^ UMI-bearing reads per cell), which is what makes the slopes comparable across rows: an absolute slope quoted without its coverage is not interpretable (§C.3).

| Library | Cells | Median UMI-reads/cell | logQ-slope (IQR) | med. $\hat{w}_{\text{tso}}$ | med. $\hat{w}_{\text{pcr}}$ |
| --- | --- | --- | --- | --- | --- |
| FLASH-UMI | 168 | 334,792 | -0.56 (-0.57, -0.54) | 0.400 | 0.005 |
| SS3:0.1uFWD | 48 | 161,112 | -0.66 (-0.68, -0.63) | 0.400 | 0.005 |
| SS3Cln:1.0uFWD | 46 | 288,040 | -0.78 (-0.80, -0.73) | 0.200 | 0.005 |
| SS3:0.5uFWD | 47 | 237,355 | -0.73 (-0.76, -0.68) | 0.200 | 0.000 |
| SS3:1.0uFWD | 48 | 233,990 | -0.74 (-0.76, -0.70) | 0.200 | 0.000 |
| SS3Cln:0.1uFWD | 48 | 314,722 | -0.82 (-0.85, -0.80) | 0.100 | 0.005 |
| SS3Cln:0.5uFWD | 48 | 266,951 | -0.75 (-0.80, -0.68) | 0.100 | 0.005 |
| Omega-seq | 337 | 949,830 | -0.83 (-0.86, -0.82) | 0.050 | 0.005 |

### B.4 Real datasets and processing

We applied the detector to 23 published single-cell RNA-seq datasets spanning full-length and droplet protocols: Smart-seq3 [9] at three forward-primer concentrations, with and without post-RT bead cleanup (ENA PRJEB35134); Smart-seq3 with molecular spikes [7]; FLASH-seq with UMIs [8]; Smart-seq-Total (GEO GSE151334); Omega-seq [6]; and 10x Genomics libraries (3*^′^* v3 and v3.1, targeted v3, and GEM-X v4 as a HEK293T:NIH-3T3 barnyard together with four healthy-donor human PBMC libraries). The full dataset list, accessions, and per-dataset estimates are given in Supplementary Table B1. For each cell we extracted clone sizes (reads per collapsed UMI) grouped by transcript position: the Smart-seq3, FLASH-seq-UMI and Omega-seq libraries were pseudoaligned with kallisto and grouped by (transcript, UMI); the molecular-spike Smart-seq3 libraries were taken from the published zUMIs/STAR alignments and grouped by (genomic position, UMI); Smart-seq-Total was aligned with HISAT2 and grouped by (gene, 3*^′^* locus, UMI); 10x libraries were taken from CellRanger output (the molecule-info file where available, otherwise the alignment restricted to uniquely mapped reads, NH=1, which removes multimapped whole-UMIs and ribosomal UMI collisions without biasing 3*^′^* counts). Cells with fewer than 10^4^ UMI-bearing reads were excluded, and the detector was run per cell (Fig. 2A).

### B.5 Downstream-damage and correction metrics

For the downstream analysis (Fig. 2B,C) we simulated paired libraries *A* and *B* of *G* = 600 genes (15,000 transcripts, *p*_rt_ = 0.6, *p*_pcr_ = 0.7, *N* = 20) at matched coverage, for a grid of (*w*_tso_*, w*_pcr_) with at least eleven replicate pairs. Writing *n*_rt*,g*_ for the true molecule count of gene *g* and *n*_det*,g*_ for its detected (phantom-inflated) UMI count, the *count inflation* is median*_g_ n*_det*,g*_*/n*_rt*,g*_ over genes with *n*_rt_ ≥ 5. Fold-change fidelity is the Spearman correlation *ρ* between the true and observed log_2_ fold-changes, 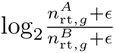 versus 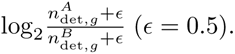 After correction (§B.6) yielding *n̂*_rt*,g*_, the residual count error is median*_g_* |*n̂*_rt*,g*_ − *n*_rt*,g*_|*/n*_rt*,g*_, and the fold-change error is (2^MAE^ − 1) × 100%, where MAE is the mean absolute deviation, in log_2_ units, between observed and true fold-changes. Points are medians over replicates and error bars are 1.4826× the median absolute deviation. Within each panel, colour encodes one rate; the solid line sets the orthogonal rate to zero and the dashed lines vary it.

**Fig. B6.**
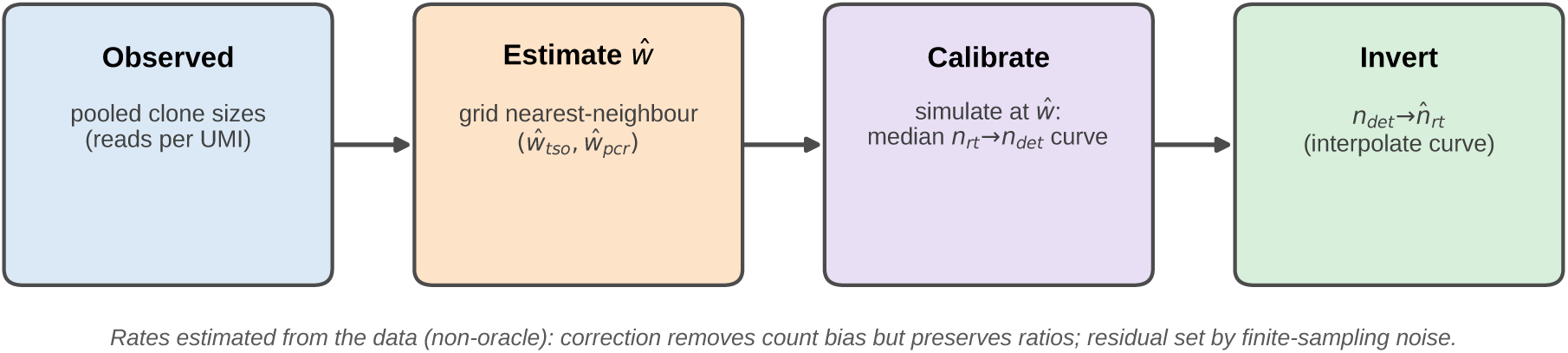
Data-estimated count-correction pipeline. Observed clone sizes → estimate (*ŵ*_tso_*, ŵ*_pcr_) by grid nearest-neighbour → simulate a calibration curve at *ŵ* → invert *n*_det_ → *n̂*_rt_.

### B.6 Count-correction pipeline

The correction is applied blind, using rates estimated from each library rather than the true (oracle) values (Fig. B6). From the pooled clone-size distribution the detector returns (*ŵ*_tso_*, ŵ*_pcr_); we then simulate libraries at those estimated rates to build the median mapping from true molecule count *n*_rt_ to detected UMI count *n*_det_, and invert this calibration curve to recover *n̂*_rt_ from each observed count. Because the mapping is a near-global rescaling, it removes the count bias while preserving count ratios (hence fold-changes are unchanged); the residual per-gene error reflects finite-sampling scatter about the median curve.

**Fig. S1.**
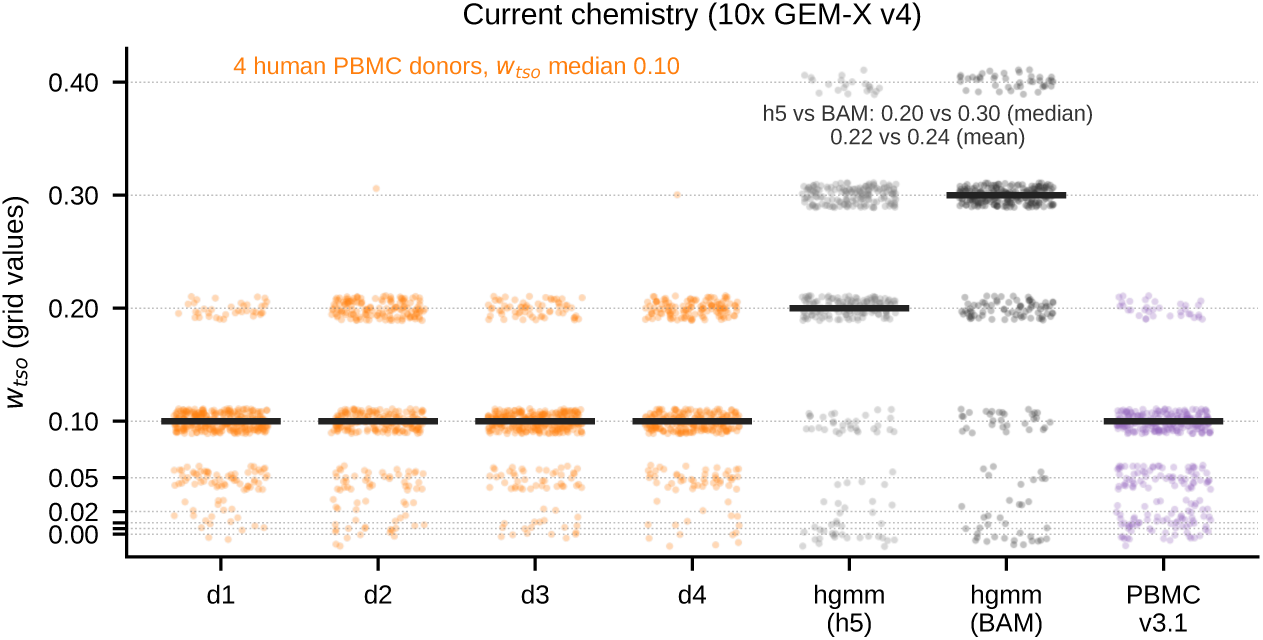
Phantom UMIs in clean human PBMC (10x GEM-X v4). Per-cell *w*_tso_ for the four healthy-donor PBMC libraries (dots, cells; bars, medians), all with median 0.10 (pooled mean 0.11). The barnyard GEM-X v4 library is scored from both its molecule-info file (h5) and its alignment (BAM), bracketing the offset between the two representations (medians 0.20 vs 0.30; means 0.22 vs 0.24), and a human-PBMC v3.1 library is shown for comparison. Every cell passing the ≥ 10^4^ UMI-read filter is included (*n* = 5,500, 5,330, 4,533, 5,144, 10,989, 5,789, 10,240 left to right). Ticks are the discrete simulation-grid values the detector matches to.

## Appendix C Supplementary information

### C.1 Phantom UMIs in current 10x GEM-X v4 chemistry

The four healthy-donor human PBMC GEM-X v4 libraries in Fig. 2A demonstrate that the artifact is present in mainstream current-generation data rather than only in specialised full-length protocols. They were quantified from the CellRanger molecule-info (h5) files, and all four centre on *w*_tso_ ≈ 0.10 (median; pooled mean 0.11) with *w*_pcr_ ≈ 0.005 (per-donor breakdown in Supplementary Fig. S1). A barnyard HEK293T:NIH-3T3 GEM-X v4 library scored from its alignment (BAM) gives a higher *w*_tso_ ≈ 0.30, at a shallower estimated coverage still (0.6 reads per true molecule, versus 1.3–1.7 for the donors); the molecule-info (h5) representation under-reports by roughly one grid step relative to the BAM, but the two agree in magnitude, and an independent human-PBMC 10x v3.1 library (*w*_tso_ ≈ 0.10) cross-confirms the donor value. Because the bead-tethered UMI-bearing oligo cannot be washed out before amplification, residual capture-primer repriming provides a plausible mechanism for this reproducible signature.

These libraries sit close to the detection floor, so the estimate should be read as a lower bound. Their estimated coverage, the reads per true molecule of the matched grid cell, is 1.7, 1.3, 1.6 and 1.6 for donors 1–4 (medians over 5,500, 5,330, 4,533 and 5,144 cells; 2.0 × 10^4^–2.3 × 10^4^ UMI-bearing reads per cell), i.e. within the coverage 1–2 regime where the clone-size distribution carries least information. (Reads per *detected* UMI is not used as a coverage measure anywhere here: its denominator is inflated by the phantoms being measured, §B.1.) In that regime the estimator is biased towards zero rather than away from it. The coverage sweep of Supplementary Fig. S4C,D begins at coverage 1, so we checked the direction of the bias at this operating point directly: simulating cells of 5 × 10^4^ true molecules with known *w*_tso_ at coverage 0.61 and re-estimating them against the same grid (24 replicates per condition; *w*_pcr_ = 0.005, *p*_pcr_ = 0.7, 12 or 20 cycles) returns medians of 0.10–0.20 for a true rate of 0.20, 0.20–0.30 for a true 0.30, and 0.30 for a true 0.40. Across all contaminated conditions only 8% of replicates exceeded their true rate, and none did at *w*_tso_ = 0.4. Clean simulations at the same coverage were called contaminated in 1.4% of replicates, whereas every one of the 5,789 barnyard cells and 99.6% of the 20,507 donor cells are called contaminated here. The quoted values are therefore floors, low by up to one grid step, and the same shallowness means they cannot be finely ranked: at this coverage a true 0.30 and a true 0.40 both return 0.30. The h5-versus-BAM offset noted above points the same way.

**Fig. S2.**
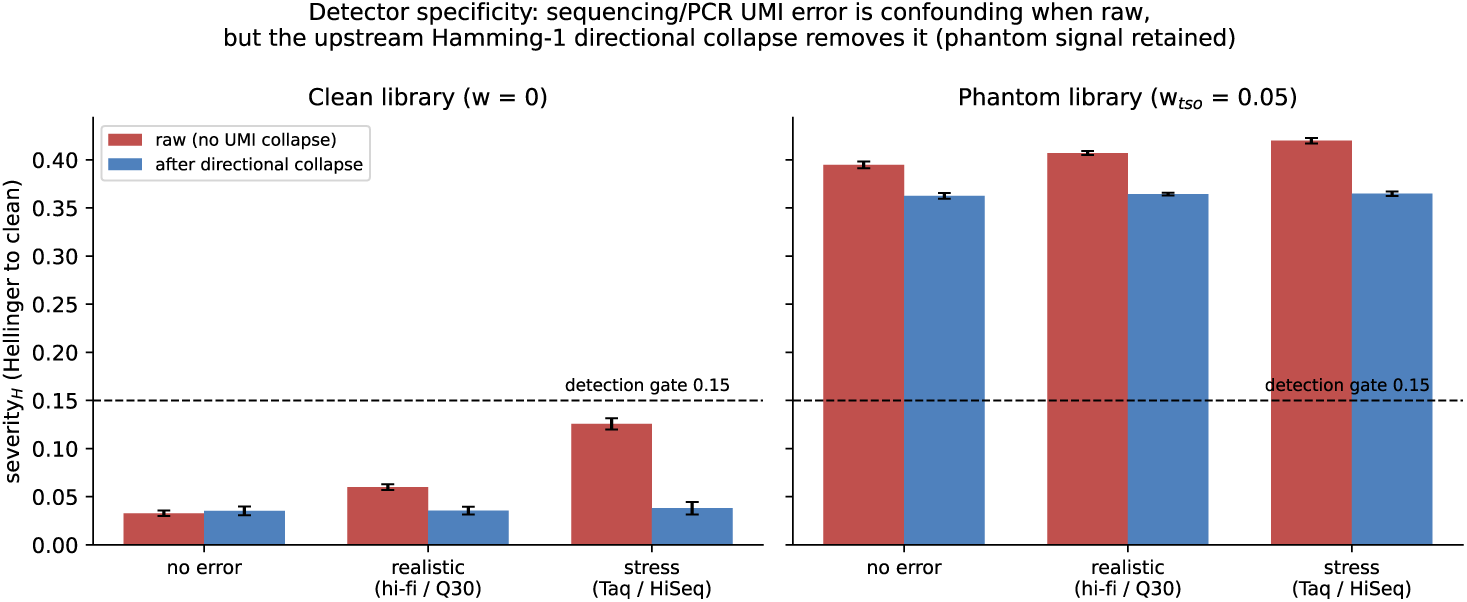
Detector specificity to UMI sequencing error. Raw severity is inflated by simulated sequencing error but returns to the clean baseline after directional UMI collapse, confirming the detector is not driven by sequencing error.

### C.2 Detector specificity to sequencing error

The clone-size signature could in principle be confounded by UMI sequencing errors, which spawn erroneous near-duplicate UMIs and add spurious small clones. On simulated libraries with realistic per-base error, the raw severity is inflated before UMI collapse, but directional (network) UMI collapse [5] removes the error-born singletons and restores severity to the clean baseline (Supplementary Fig. S2). The detector therefore responds to amplification-born phantom UMIs, not to sequencing error, consistent with the mechanistic distinction drawn in the main text.

### C.3 A single index is insufficient for parameter-blind detection

A single summary statistic such as the logQ-slope detects contamination well when the coverage and PCR efficiency are known: per-coverage detection AUC is near unity, at low coverage even exceeding that of the severity*_H_* statistic. Its clean baseline, however, is not a fixed constant: it drifts from near −1 at deep coverage toward ≈ −0.7 when reads per UMI are few (Supplementary Fig. S3A), overlapping the contaminated range. Blind to coverage and efficiency, no single global threshold separates clean from contaminated libraries; a logQ-slope cutoff of −0.95 (tuned for deep clean data) misclassifies 53% of clean libraries, and 100% of the shallowest. The severity*_H_* statistic, referenced to the clean grid at each library’s own coverage and efficiency, keeps a flat clean baseline (≈ 0.03; Supplementary Fig. S3B). Consequently the parameter-blind detection AUC of severity*_H_* (0.72–0.98) exceeds that of the logQ-slope (0.62–0.96) at every contamination level (Supplementary Fig. S3C). This motivates detecting phantoms from the full binned clone-size distribution rather than a single scalar. The drifting baseline is a statement about *parameter-blind, absolute* thresholds, and it does not apply to the way the logQ-slope is used in the companion study [6], where libraries prepared and sequenced side by side are compared with one another at matched coverage: a relative comparison cancels the coverage dependence that defeats a global cutoff. The distinction matters in practice, since the clean baseline at low coverage (≈ −0.7) approaches the slopes reported there for phantom-free libraries (≈ −0.8), so a slope quoted without its coverage is not interpretable on its own. Used that way the two metrics agree: across the eight deep full-length libraries scored by both studies (Smart-seq3 at three forward-primer concentrations ± pre-PCR cleanup, FLASH-seq-UMI and Omega-seq), the per-library logQ-slope and *ŵ*_tso_ rank the libraries almost identically (Spearman *ρ* = 0.90; Supplementary Table B3), with Omega-seq steepest and cleanest and FLASH-seq-UMI shallowest and dirtiest at both ends.

### C.4 Coverage and PCR-efficiency sweeps

Sweeping coverage and PCR efficiency *p*_pcr_ (Supplementary Fig. S4) shows that: (i) detection AUC rises with coverage and, for *w*_tso_, is reached at slightly higher coverage as *p*_pcr_ increases, while *w*_pcr_ is detected by coverage ≈ 3 regardless of efficiency (panels A,B); (ii) the detector’s estimates *ŵ*_tso_ and *ŵ*_pcr_ under-shoot at low coverage but converge to the true rates by coverage ≈ 8–13 (panels C,D); and (iii) at fixed coverage the normalized clone-size severity collapses onto *w*_tso_ (panel E; the *p*_pcr_ curves overlap) but not onto the product *a* = *p*_pcr_*w*_tso_ (panel F). The repriming rate *w*_tso_ is thus directly identifiable from the shape of the distribution, with *p*_pcr_ acting only as a weak amplification scale that largely cancels in the normalized histogram.

**Fig. S3.**
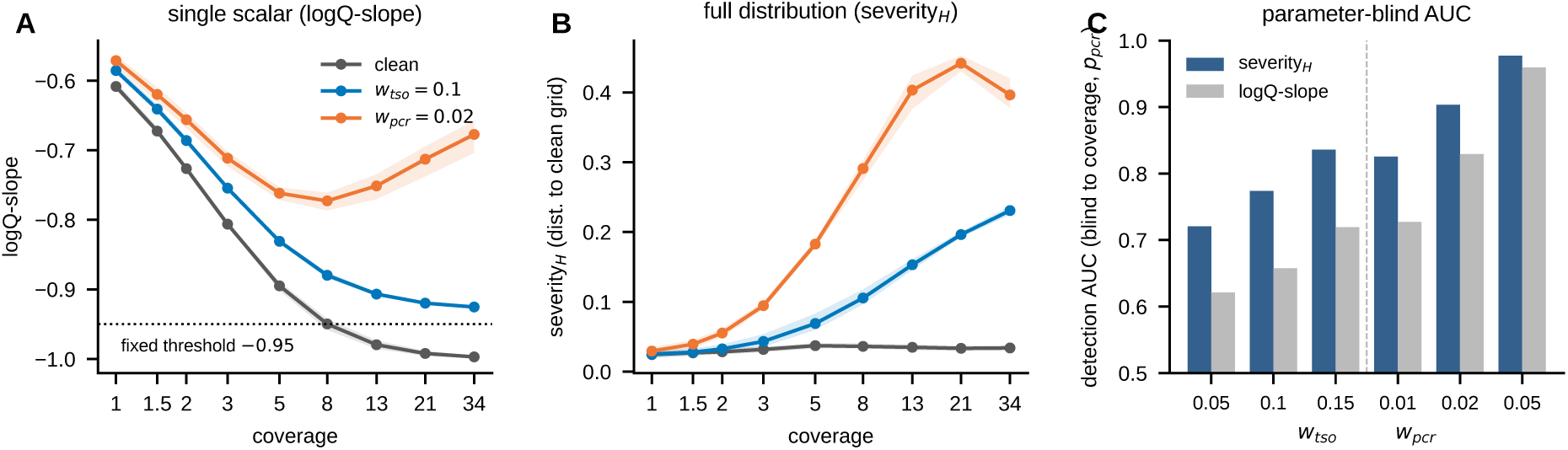
Single-index insufficiency. **A**, logQ-slope versus coverage: the clean baseline (grey) drifts into the contaminated range; a fixed threshold (dotted) misclassifies shallow clean libraries. **B**, severity*_H_* versus coverage: the clean baseline is flat (grid-referenced). **C**, parameter-blind detection AUC (single global threshold; coverage, *p*_pcr_ unknown): severity*_H_* exceeds logQ-slope at every level.

**Fig. S4.**
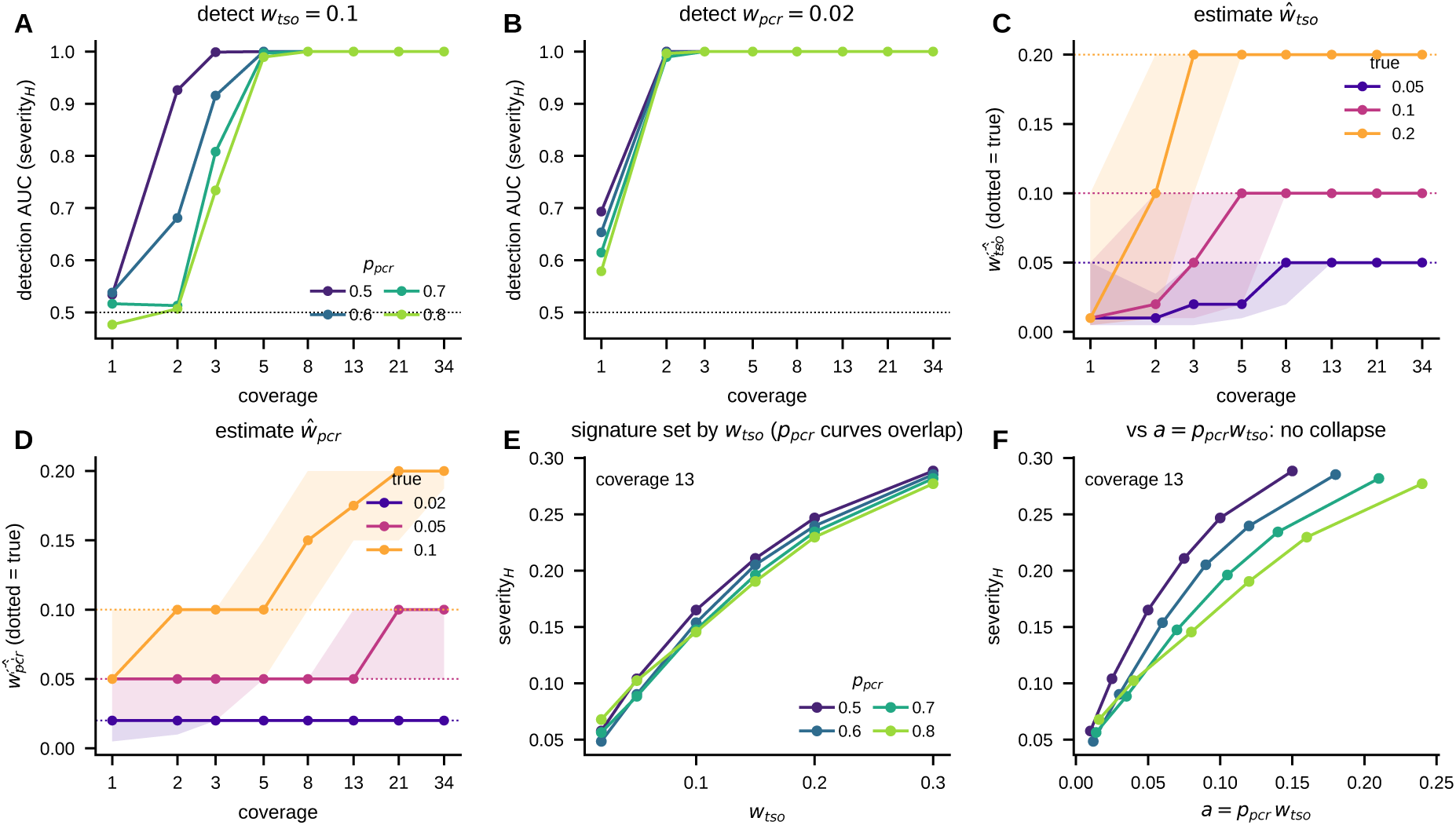
Coverage and PCR-efficiency sweeps. **A,B**, detection AUC versus coverage, split by *p*_pcr_, for *w*_tso_ and *w*_pcr_. **C,D**, detector estimates *ŵ*_tso_, *ŵ*_pcr_ versus coverage (median ± IQR; dotted, truth). **E,F**, at coverage 13, severity*_H_* collapses on *w*_tso_ (E) but not on the rate *a* = *p*_pcr_*w*_tso_ (F).

**Fig. S5.**
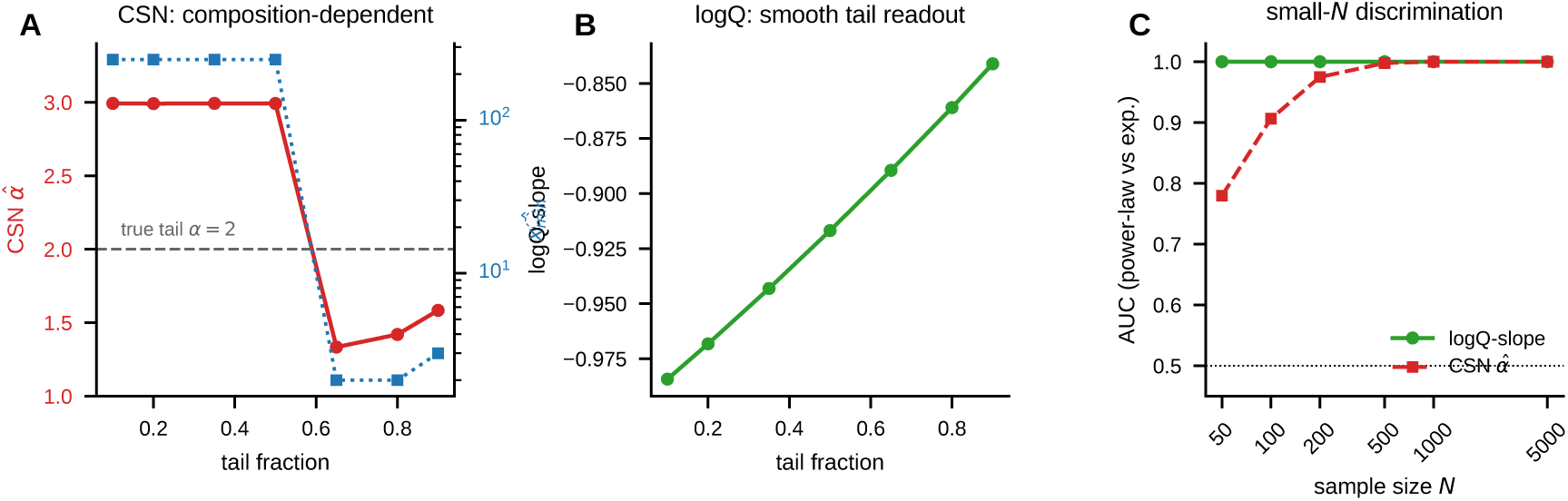
logQ-slope versus Clauset–Shalizi–Newman power-law fitting. **A**, On a uniform-plus-power-law mixture (true tail *α* = 2, dashed), CSN *α̂* (red) and *x̂*_min_ (blue) are set by the tail fraction, not the tail exponent, and jump discontinuously. **B**, the logQ-slope varies smoothly with tail fraction. **C**, AUC for discriminating a power-law from an exponential tail versus sample size *N* : the logQ-slope separates them at *N* = 50, whereas CSN *α̂* requires *N* >̰ 500.

### C.5 logQ-slope versus standard power-law fitting

The power-law exponent *β* of the phantom clone-size distribution (Methods) could in principle be estimated directly from data by the standard Clauset–Shalizi–Newman (CSN) procedure, which selects a lower cutoff *x*_min_ by Kolmogorov–Smirnov minimisation and fits the exponent *α̂* by maximum likelihood above it. On the mixture distributions that phantom clone-size data present (a bulk of true UMIs plus a power-law tail of phantoms), this fails: *α̂* and *x̂*_min_ are dictated by the bulk-to-tail composition rather than the tail exponent, jumping discontinuously as the tail fraction grows and never recovering the true tail *α* (Supplementary Fig. S5A). The logQ-slope, which needs no *x*_min_, instead varies smoothly and monotonically with tail heaviness (Supplementary Fig. S5B), and discriminates a power-law from an exponential tail even at *N* ≈ 50 observations, where CSN requires *N* >̰ 500 (Supplementary Fig. S5C). This supports detecting phantoms from the full binned clone-size distribution and, as a scalar summary, the logQ-slope rather than a fitted power-law exponent.

